# Excitation/Inhibition balance relates to cognitive function and gene expression in Temporal Lobe Epilepsy: an hdEEG assessment with aperiodic exponent

**DOI:** 10.1101/2024.02.01.578441

**Authors:** Gian Marco Duma, Simone Cuozzo, Luc Wilson, Alberto Danieli, Paolo Bonanni, Giovanni Pellegrino

## Abstract

Patients with epilepsy are characterized by a dysregulation of excitation-inhibition balance (E/I). The assessment of E/I may inform clinicians during the diagnosis and therapy management, even though it is rarely performed. An accessible measure of the E/I of the brain represents a clinically relevant feature. Here we exploited the exponent of the aperiodic component of the power spectrum of EEG signal as a noninvasive and cost-effective proxy of the E/I balance. We recorded resting-state activity with high-density EEG from 65 patients with temporal lobe epilepsy (TLE) and 35 controls. We extracted the exponent of the aperiodic fit of the power spectrum from source-reconstructed EEG and tested differences between TLE and controls. Spearman’s correlation was performed between the exponent and clinical variables (age of onset, epilepsy duration and neuropsychology) and cortical expression of epilepsy-related genes derived from Human Allen Brain Atlas. Patients with TLE showed a significantly larger exponent, corresponding to an inhibition directed E/I balance, in bilateral frontal and temporal regions. Lower E/I in the left entorhinal, and bilateral dorsolateral prefrontal cortices corresponded to a lower performance of short term verbal memory. Limited to TLE, we detected a significant correlation between the exponent and the cortical expression of GABRA1, GRIN2A, GABRD, GABRG2, KCNA2and PDYN. EEG aperiodic exponent maps the E/I balance non-invasively in patients with epilepsy and reveals a tight relationship between altered E/I patterns, cognition and genetics.

## 1 Introduction

Epilepsy is a disorder characterized by a dysregulation of the excitation/inhibition balance (E/I) as suggested by converging evidence ranging from genetics, to proteomics expression, to clinical presentation ^1–4^. Altered E/I balance translates into an increased risk of recurrent epilepsy seizures ^5–7^. Hence, pharmacological treatments aim to reduce the risk of seizures by interfering with E/I through several mechanisms of action ^8–10^. Despite its biological and clinical relevance, E/I balance is rarely directly investigated in epilepsy, most likely because its estimation is difficult and time-consuming. Fortunately, recent developments offer a streamlined assessment of E/I balance, leveraging properties of spontaneous electromagnetic signals which can be easily measured with magneto- or electroencephalography at high spatial and temporal resolution, providing extensive brain coverage with potential translations in a clinical setting ^11–14,15(p2)^. Amongst multiple signal features, the exponent of the aperiodic component of the M/EEG signal power spectrum provides information related to system E/I balance ^16–18^.

In this study, we investigated E/I balance in patients with temporal lobe epilepsy (TLE) in comparison to healthy subjects, by measuring the aperiodic exponent obtained from cortical reconstructed high-density EEG signals at rest. From a clinical perspective, TLE is of particular interest as it is one of the most prevalent and homogeneous manifestations of focal epilepsy ^19,20^. Considering the network nature of this neurological disorder, we hypothesized to detect more prominent E/I alterations in brain regions related to seizure onset, namely temporal lobe, as well as interconnected regions. Moreover, we expected to detect patterns of E/I changes influenced by the laterality of the epileptic focus. We further hypothesized a relation between E/I balance and clinical variables as well as an influence of antiseizure medications due to their known interaction with this property ^21,22^. Disrupted E/I balance may impact synaptic plasticity which represents one of the scaffolding mechanisms allowing a system to efficiently respond to environmental stimuli ^23,24^. Therefore, altered E/I balance may be connected to the impairments detected in the TLE population in different cognitive domains and linked to functional network disruption at multiple spatial scales ^25–28^. As such, we would expect to detect a relationship between the aperiodic exponent and the neuropsychological profiles in patients with TLE. Importantly, E/I balance is a multifactorial byproduct of neuronal activity, influenced by environmental factors as well as gene expression of channels and receptors ^29,30^. Numerous mutations of gene regulating gamma-aminobutyric acid receptors, as well as sodium-potassium channels have been enlisted as etiopathogenic causes of epilepsy ^31–34^. The recent release of open transcriptomics datasets such as the Allen Human Brain Atlas (AHBA) has offered opportunities to explore how gene expression patterns in the brain reflect macroscale neurofunctional findings ^35,36^. Functional-genetic information integration can shed light on the micro- to macroscale pathophysiological mechanisms underlying epilepsy. Here, we exploited transcriptomic data in order to connect epilepsy-related genes with E/I balance. Overall, the present study investigates the feasibility of the aperiodic component of the PSD via hdEEG as a noninvasive measure of E/I balance, linking it to clinical, neuropsychological, and genetic factors to elucidate the neural mechanisms underlying brain dynamics and cognitive alteration in TLE.

## 2 Method

### 2.1 Participants

We enrolled patients with temporal lobe epilepsy who underwent high-density electroencephalography (hdEEG) for clinical evaluation between 2018-2022 at the Epilepsy and Clinical Neurophysiology Unit, IRCCS Eugenio Medea in Conegliano (Italy) and a group of healthy controls. The diagnosis of temporal lobe epilepsy was established according to the ILAE guidelines. The clinical workflow included Video EEG (32 channels) monitoring, brain magnetic resonance imaging (MRI), and positron emission tomography (PET) as an adjunctive investigation in selected patients. Seventy-two cases were retrospectively screened, 5 patients were excluded due to previous brain surgery or invasive investigation, resulting in a final sample size of 67 (mean age = 37.18 [SD = 18.04]; 24Lfemales; 30 left-TLE; 17 right-TLE; 20 bilateral TLE). Patients’ demographic and clinical characteristics are provided in Table 1. The control group was composed of 35 healthy participants with no prior history of neurological or psychiatric disorders (mean age = 34.92 [SD = 9.22]; 25Lfemales). The study protocol was conducted according to the Declaration of Helsinki and approved by the local ethical committee. All participants signed a written informed consent.

**Table 1.**
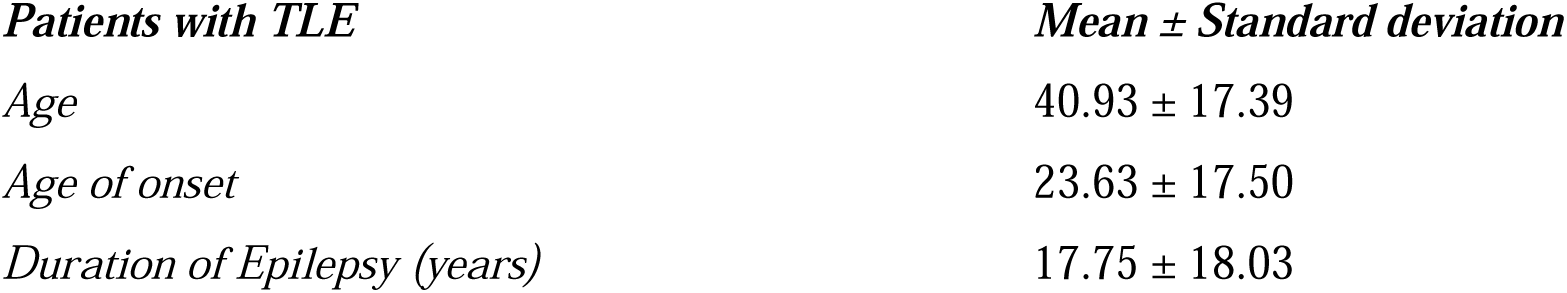
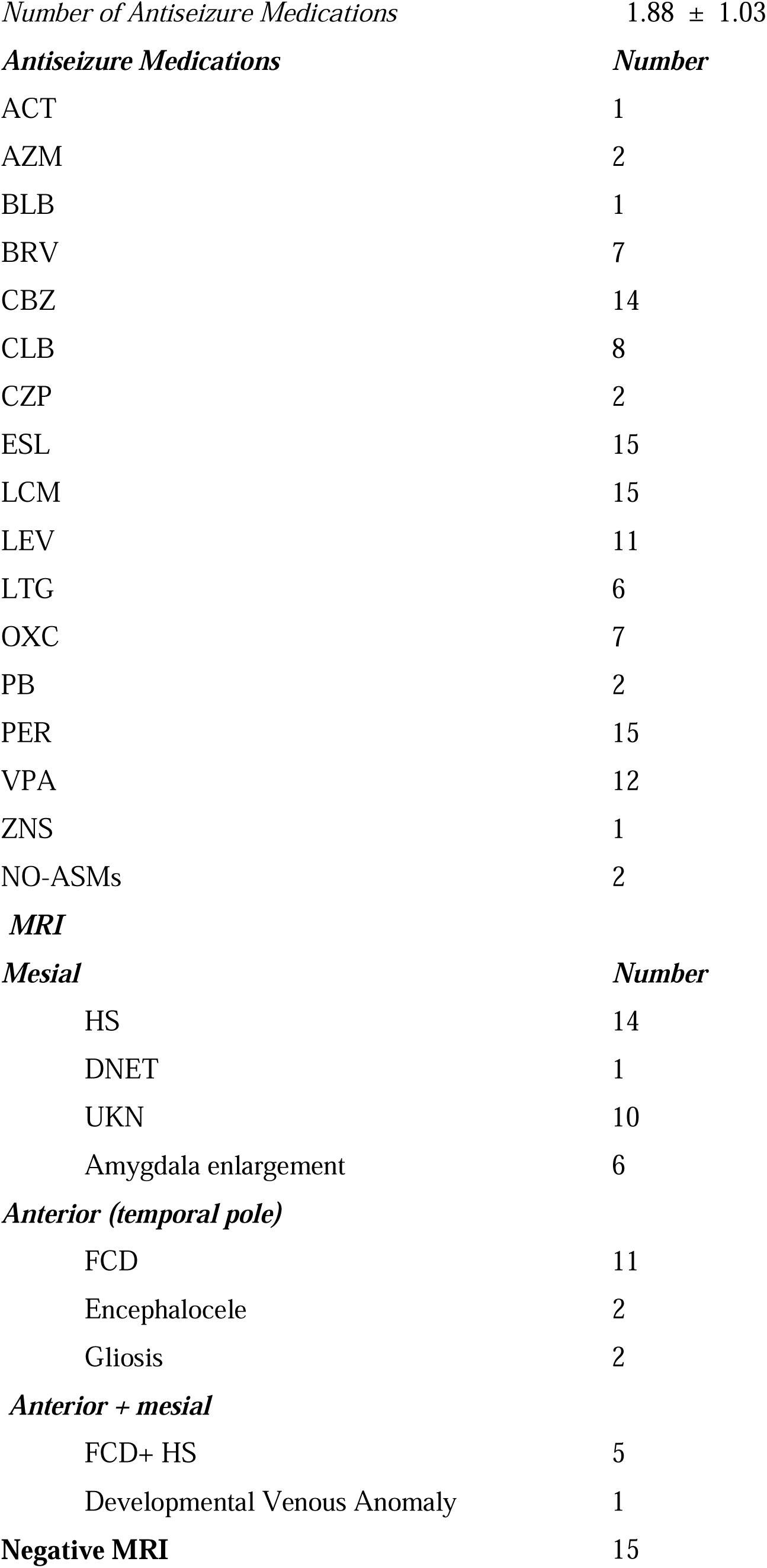
Sample demographic and clinical information. Demographic and clinical features of the patients. MRI findings are reported by sublobar localization. The continuous variables are reported as mean ± SD. Antiseizure medications abbreviations: ACT, acetazolamide; AZM, acetazolamide; BRV, brivaracetam; CBZ, carbamazepine; CLB, clobazam; CZP, clonazepam; ESL, eslicarbazepine; LCM, lacosamide; LEV, levetiracetam; LTG, lamotrigine; OXC, oxcarbazepine; PB, phenobarbital; PER, perampanel; VPA, valproic acid; ZNS, zonisamide; NO-ASMs, no pharmacological treatment. Abbreviation MRI abnormalities: FCD, focal cortical dysplasia; HS, hippocampal sclerosis; DNET, dysembryoplastic neuroepithelial tumours; UKN, unknown.

### 2.2 Neuropsychological assessment

Neuropsychological assessment focused on memory, attention/executive functions and intelligence. Short-term memory and long-term memory functioning (STM) were assessed using the Digit Span ^37^, Corsi block tapping tests ^37^,Rey–Osterrieth Complex Figure Test (ROCFT) ^38^ and Rey Auditory Verbal Learning Test ^39^. Attention and executive functions were evaluated using the Trail Making Test (TMT) (part A and B) ^40^. Finally, we used the total IQ of the WAIS-IV ^41^ or WISC-IV ^42^ scales as a measure of global intelligence. Table 2 shows the descriptive statistics of the neuropsychological scores.

**Table 2.** Neuropsychological scores. Abbreviation: IQ = intelligent quotient; RAVLT = Rey auditory verbal learning test; ROCF = Rey–Osterrieth complex figure test; TMT A/B = trail making test A/B

| Test | Score (mean ± standard dev) |
| --- | --- |
| *Digit Span* | 5.73 ± 1.06 |
| *Corsi block Tapping Test* | 4.90 ± 1.11 |
| *ROCFT - Copy* | 30.81 ± 7.20 |
| *ROCFT - Reproduction* | 15.03 ± 6.22 |
| *RAVLT - Immediate* | 39.01 ± 9.75 |
| *RAVLT - Delayed* | 7.17 ± 3.24 |
| *Total IQ* | 90.13 ± 19.62 |
| ***TMT-A*** | 34.47 ± 18.05 |
| ***TMT-B*** | 110.25 ± 76.84 |

### 2.3 Resting State EEG recording

Ten minutes of eyes-closed resting state hdEEG was recorded with a 128-channel Micromed system while participants sat comfortably on a chair in a silent room. The signal was sampled at 1,024 Hz and referenced to the vertex. The impedance of the electrodes was kept < 5kΩ, for each sensor.

### 2.4 EEG pre-processing

Signal preprocessing was performed via EEGLAB 14.1.2b ^43^ according to a pipeline validated in our previous work ^26,44^. The preprocessing pipeline included the following steps: a) resampling to 250 Hz; b) bandpass-filter (0.1 to 45 Hz) with a Hamming windowed sinc finite impulse response filter (filter order = 8250); c) epoching (1 second-long epochs); d) removal of epochs and channels containing IEDs and artifacts; e) removal of flat channels; f) independent component analysis ^45^ (Stone, 2002), using the Infomax algorithm ^46^ as implemented in EEGLAB; g) channel interpolation with spherical spline interpolation method ^47^; h) re-reference to average.

In further details, interictal epileptiform discharges (IEDs) were identified by trained personnel (GMD, AD and PB). Epochs containing IEDs were purposely removed to assess intrinsic brain functional organization independently from epileptiform activity; automated bad-channel and artifact detection and removal was performed with the TBT plugin implemented in EEGLAB, with the following parameters: channels differential average amplitude 250μV in more than 30% of the epochs. Epochs were removed if they contained IEDs, or included more than 10 bad channels. Flat channels were identified and removed with the Trimoutlier EEGLAB plug-in setting a within the lower bound of 1μV. We rejected an average of 46.80 ± 44.40 (SD) epochs and an average of 9.51± 4.68 (SD) components. After preprocessing, each subject had at least 6 minutes of artifact-free signal.

### 2.5 Cortical Source modeling

For most of the TLE patients, the individual anatomic MRI for source imaging consisted of a T1 isotropic three-dimensional (3D) acquisition. For twelve patients and for the control group we used the MNI-ICBM152 default anatomy ^48^. The MRI was processed with the Computational Anatomy Toolbox (CAT12) ^49^, to obtain skin, skull, and gray matter surfaces. Anatomical data was then imported in Brainstorm ^50^. The co-registration of the EEG electrodes with brain MRI was performed using Brainstorm, considering anatomical landmarks. Manual coregistration corrections were applied as needed. The cortical mesh was downsampled at 15,002 vertices and the three Boundary Element Models (BEM) surfaces were reconstructed (inner skull, outer skull and head). We computed the forward model with the 3-shell BEM approach, setting the tissue connectivity of brain, skull and skin to 0.33, 0.165, 0.33 S/m; (ratio: 1/20), respectively. This was achieved with the OpenMEEG plugin ^51,52^ implemented in Brainstorm. We used the weighted minimum norm as inverse solution method, with Brainstorm’s default parameters settings ^53^.

### 2.6 Parametrizing brain spectra

Prior to performing the parametrization of brain spectral components, source activity was downsampled to 68 cortical regions of interest (ROIs) according to the Desikan-Killiany atlas^54^. ROI time-series were obtained as the mean time series across ROI vertices ^55^. For each ROI time series, the power spectrum was calculated using Welch’s method (Hann window, length 5 seconds, overlap 50%). Python “fooof’ library was then used in order to parameterize the neural power spectrum into its periodic and aperiodic components ^16^. The exponent X of 1/f^X^-like aperiodic activity was calculated between 1-35 Hz. Fitting was performed using the ‘fixed’ aperiodic mode. Spectral parameterization settings for the algorithm were: peak width limits =L[0.5,12], maximum number of peaks = inf, peak threshold =L2, minimum peak height =L0.0.

### 2.7 Cortical gene expression from the Allen Human Brain Atlas

Regional microarray expression data were obtained from six post-mortem brains of the Allen Human Brain Atlas (AHBA; http://human.brain-map.org/) ^56^. We used the abagen toolbox (https://github.com/netneurolab/abagen; ^35^ to process and map the data to the 68 ROIs of the Desikan-Killiany parcellation ^54^. To begin, the microarray probes were reannotated using information provided by Arnatkeviciute and colleagues ^57^. The reannotated information discarded probes without a reliable match to genes. Probes were then filtered based on their expression intensity relative to background noise ^58^. Specifically, probes with an intensity less than background in ≥50% of samples were discarded. When multiple probes indexed the expression of the same gene, the probe with the most consistent pattern of regional variation (i.e., differential stability; ^56^ across donors was selected. Differential stability was calculated as it follows:

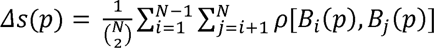

where ρ is Spearman’s rank correlation of the expression of a single probe *p* across regions in two donor brains B*i* and B*j*, and N is the total number of donors. This procedure retained 15,656 probes, each representing a unique gene.

Samples were assigned to brain regions in the provided atlas if their MNI coordinates were within 2 mm of a given parcel. To reduce the potential for misassignment, sample-to-region matching was constrained by hemisphere and cortical/subcortical divisions ^57^. All tissue samples not assigned to a brain region in the provided atlas were discarded.

Inter-subject variation was addressed by normalizing tissue sample expression values across genes using a robust sigmoid function ^59^:

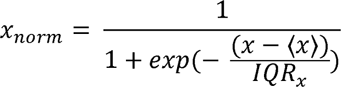

where (x) is the median and IQR_x_ is the normalized interquartile range of the expression of a single tissue sample across genes. Normalized expression values were then rescaled to the unit interval.

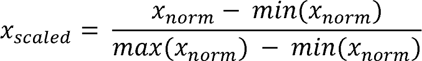

Gene expression values were then normalized separately for each donor across regions using an identical procedure. Scaled expression profiles were finally averaged across donors, resulting in a single matrix with rows corresponding to brain regions and columns corresponding to the retained 15,656 genes.

Based on a recent panel of epilepsy-related genes ^60^ we extracted a set of 22 genes linked with the regulation of intra and intercellular E/I balance. These selected genes regulate the functioning of sodium-potassium channels, gamma-aminobutyric acid (GABA) and N-methyl-D-aspartate (NMDA) receptors, seizure susceptibility and the presence of focal cortical dysplasia. A detailed list of the included genes and the associated functions is provided in the supplementary material (see Supplementary Table 1).

### 2.8 Statistics

We first contrasted the exponent value of all the patients with epilepsy vs. controls, for each of the ROIs, using an independent samples two-tailed *t*-test with 5000 permutations. We then repeated the same analysis contrasting specific patient groups against healthy controls, namely Left-TLE, Right-TLE and bilateral-TLE. Correlations between the exponent and clinical (age of onset, epilepsy duration and number of ASMs), neuropsychological (test scores), as well as gene expression variables were assessed with Spearman’s correlation coefficients. Notably, exponent-gene correlation was performed in both TLE patients and controls, to highlight specific patterns related to epilepsy. The significance level was set to *p* < 0.05. The alpha inflation due to multiple comparisons was controlled with the false discovery rate (FDR) procedure ^61^, across ROIs (N=68) or genes (N=22), as appropriate. Brain plots were obtained using the enigma-toolbox ^62^.

## 3 Results

### 3.1 Exponent differences in TLE vs. Controls

We observed that patients with TLE are characterized by a larger aperiodic exponent (i.e., steeper spectral slope, lower E/I) as compared to healthy controls. Significant differences were mainly observed in regions involved in seizure propagation such as superior and inferior temporal gyri, entorhinal cortex and temporal pole, anterior cingulate cortex and in the posterior quadrant (cuneus and visual cortex) (all *p*_adjusted_ < .05; see Fig2. A). When considering the three groups separately, patients with both right and left TLE showed a larger exponent in the left temporal areas (all *p*_adjusted_ < .05), whereas bitemporal epilepsy (BTLE) showed an increase in both temporal and frontal regions, bilaterally (all *p*_adjusted_ < .05). All regional statistical values (*t-value* and the corresponding *p-value* for each significant ROI) are provided in the supplementary materials (Supplementary Table 2-5).

### 3.2 Clinical variable and neuropsychology correlations

A significant positive relationship between the aperiodic exponent and the number of ASMs was found in the entire group of patients, suggesting that a larger number of antiseizure medications corresponded to lower E/I. The correlation was spatially expressed in the medial temporal regions as well as in the frontal gyrus and dorsolateral prefrontal cortex (DLPFC) (*rho_max_* = .41; *p_adjusted_* < .05; see Fig.3A). A significant negative correlation was found between aperiodic exponent and short term verbal memory (RAVLT) in the left temporal regions, the DLPFC, and cuneus bilaterally, with larger exponent (lower E/I) linked to a lower cognitive performance (*rho_max_* = - .49; *p_adjusted_*< .05; see Figure.3A). All regional statistical values (*Spearman’s rho* and the corresponding *p-value* for each significant ROI) are provided in the supplementary materials (Supplementary Table 6-7)

### 3.3 Exponent-gene correlations results

The correlation between the exponent and the gene’s cortical expression displayed a specific pattern differentiating TLE patients and controls. Specifically we observed a higher cortical expression of the genes related to the functioning of sodium channels (KCNA2; *rho* = - .395; *p_adjusted_* = .003) and with gaba receptors (GRIN2A; *rho* = - .394; *p_adjusted_* = .003; GABRA1; *rho* = - .433; *p_adjusted_*= .001; GABRD; *rho* = - .403; *p_adjusted_*= .001; GABRG2; *rho* = - .475; *p_adjusted_* < .001) associated with a larger exponent value. Additionally, we observed an opposite pattern in a gene involved in the self-regulation of the seizure through the peptide ligands of opioid receptors (PDYN; r*ho* = - .403; *p_adjusted_* = .001). By contrast, we observed no such differences in the control group.

## 4 Discussion

We exploited the exponent of the aperiodic component of the power spectrum from source derived hdEEG signals, as a non-invasive and cost-efficient proxy of the excitation/inhibition alterations in patients with epilepsy. As first we observed that patients with temporal lobe epilepsy (TLE) were characterized by larger aperiodic exponent values as compared to healthy controls. The statistical difference was maximally expressed in temporal regions, as well as dorsal frontal areas, the cingulate cortex and the precuneus (Figure 2A). This E/I imbalance displayed sensitivity to the lateralization of the focus with a left dominant exponent increase in left-TLE and a bilateral distribution in BTLE. Converging evidence suggests that higher values of the aperiodic exponent are indicators of system E/I balance shifted towards inhibition ^16, 17^. In this light, our result would suggest that patients with TLE display a more predominant inhibitory component, as assessed from non-invasive and perturbation-free methods. Our cohort includes medically refractory epilepsy patients, treated with antiseizure medications. A significant link between the number of medications and E/I shift towards inhibition was indeed observed. Therefore, it is conceivable that E/I dysregulation was at least partially counterbalanced by antiseizure medications in these patients. Importantly, the spatial configuration of the relationship suggests that fronto-temporal areas were the most affected. Multiple interpretations can account for the exponent-pharmacology relationship. Firstly, converging evidence from neuroimaging studies have shown that medial prefrontal and temporoparietal cortices display longer intrinsic neuronal time scales (INT) ^63,64^. INTs can be interpreted as the degree to which spontaneous brain activity correlates with itself over time, setting the duration over which brain regions integrate inputs ^65^. Patients with TLE are characterized by longer INTs in the temporal areas^66^. Relevantly, recent findings suggest that INTs have a direct link with the decay of the PSD and therefore the exponent value ^67^. In this light the regions characterized by slower intrinsic timescales could be the most sensitive to the ASMs, resulting in an increase of low-frequency power and a larger exponent. Secondly, homeostatic plasticity mechanisms would explain this spatial distribution. Homeostatic plasticity enables the tuning of synaptic activity in relation to environmental stimuli, in which the magnitude and direction of synaptic plasticity are adjusted according to the recent history of postsynaptic activity ^68,69^. Regions belonging to the epileptic network have an excitatory driven balance ^70^, that would be more sensitive to the effect of antiseizure medications, up to reversing their pattern ^71^. In other words, the mesial temporal lobe structures and the regions connected to them are potentially characterized by an intrinsically higher E/I ^70^. The medication effects may induce a plasticity shift towards inhibition on the E/I balance of these regions ^72^. This interpretation would be consistent with previous transcranial magnetic stimulation studies, which have detected homeostatic plasticity effects in TLE ^69,73,74^. Dysregulation of E/I balance has been proposed as one of the neurophysiological mechanisms explaining performance reduction of cognitive functioning ^75^ and may represent the mechanism explaining the observed correlation between the exponent and the memory functioning. This relationship displayed a spatial configuration involving the left temporal and bilateral dorsolateral prefrontal cortices. Neuroimaging studies identified left temporal areas as the scaffolding regions of the verbal memory ^76–78^. However, the information retention and recall engage a distributed network also involving fronto-parietal regions ^79–82^. In light of this, our findings suggest that altered E/I balance in the distributed network underlying memory may be one of the neurophysiological mechanisms explaining impairments of this cognitive domain characterizing patients with TLE ^83–85^. The response properties of neurons are indeed shaped by the balance between co-activated inhibitory and excitatory synaptic inputs ^86^. Regulation of E/I balance is a multifactorial process related to neuro-biophysical properties and dynamics mediated by neurotransmission and channel activity ^87,88^. Computational and biophysical models show that GABA and NMDA receptors as well as sodium-potassium channels are the main regulators of E/I at multiple scales, from single cell to vast neuronal assemblies ^17,89–92^. By integrating source EEG functional and transcriptional atlas data, we tested the hypothesis that transcriptomic could covary with E/I, providing information of the potential mechanisms underlying E/I deregulation in TLE. We observed a correlation of the noninvasive measure of E/I balance and the expression of the gene regulating GABA receptors and potassium channels, exclusively in the patients group. Specifically, an increased cortical expression of GABRA1, GRIN2A, GABRD and GABRG2 are related with a lower exponent. This correlational pattern is also observed in the gene regulating potassium voltage-gated channel subfamily A member 2, namely the KCNA2. Noticeably, alterations of genes’ expression regulating GABA units, as well as potassium channels, have been linked with the disruption of inhibitory network development and/or activity modulation, being considered as one of pathophysiological mechanisms underlying multiple types of epilepsy ^93–95^. Additionally, we also observed a relationship between the exponent and the cortical expression of the PDYN in the TLE patients group. This gene is involved in the synaptic plasticity via kappa-opioid receptors ^96,97^ (Schwarzer 2009; Chavkin 2013) and encodes for the anticonvulsant peptide dynorphin, a strong candidate for a seizure suppressor gene and thus a possible modulator of susceptibility to TLE ^98^ . Notably, the correlation patterns between aperiodic exponent and gene expression were exclusively identified in the clinical group. Using normative neurotransmitter atlas data to contextualize functional alterations in epilepsy has inherent limitations. The gene expression atlases were derived from an independent sample of individuals. However, these data represent a methodological unicum of their kind being a useful tool to understand the link between micro to macro-scale levels of network functions. In fact, this type of data has been successfully used in recent studies investigating the relationship between gene expression and network functioning in neurological disorders ^99–101^.

**Figure 1.**
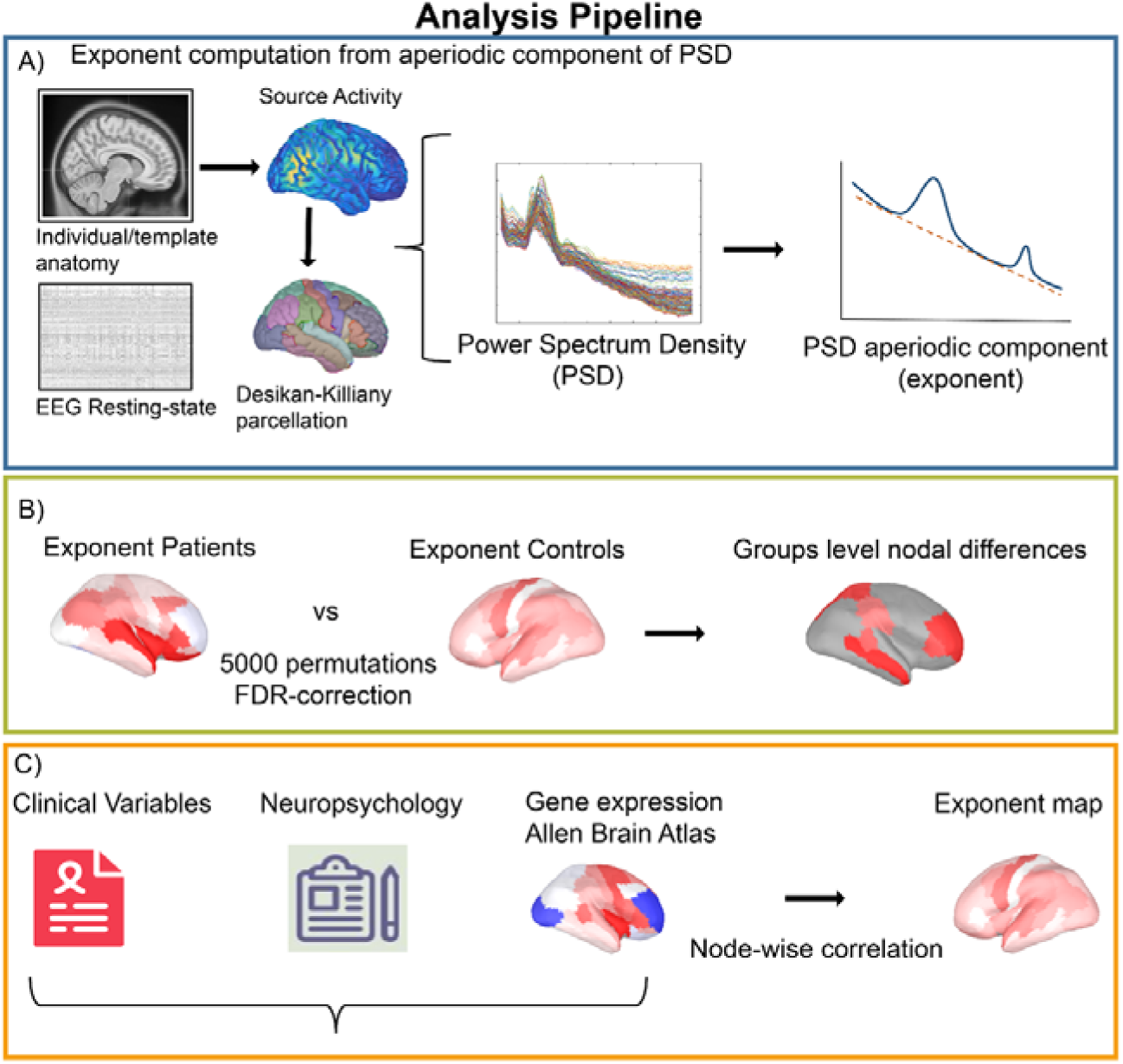
Analytical pipeline. The present figure displays the analytical steps performed. Panel A follows the steps for extracting aperiodic exponents from resting-state EEG recordings. Panel B graphically explains the statistical comparison performed. Panel C represents the correlational analysis between clinical variables, neuropsychological assessment, gene expression and aperiodic exponent.

**Figure 2.**
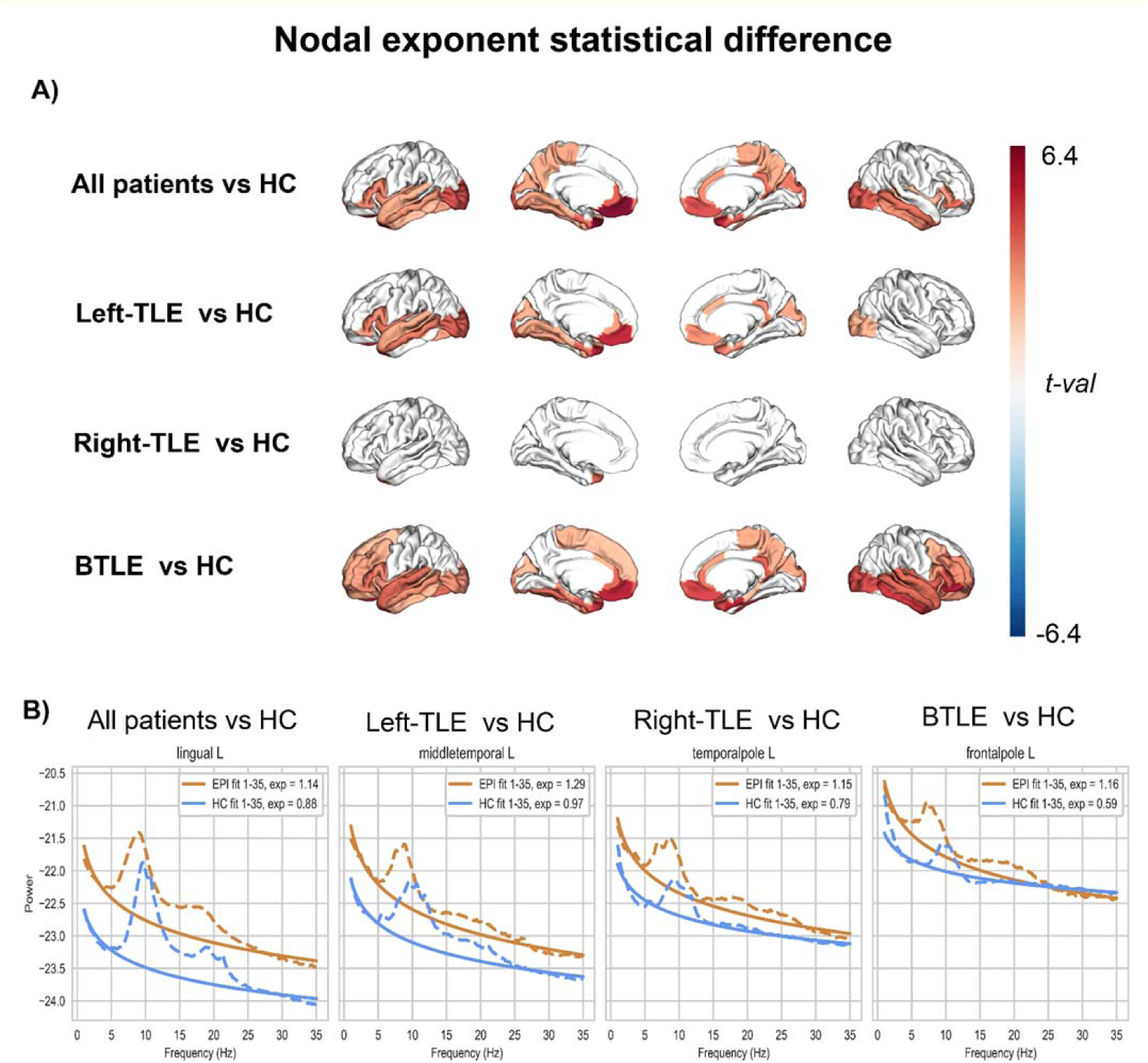
Statistical difference across groups of the nodal aperiodic exponent. Panel A displays the statistical difference of the aperiodic exponent comparing all patients with temporal lobe epilepsy (TLE), Left-TLE, Right-TLE and bilateral TLE (BTLE) vs. healthy controls. Panel B shows the power spectrum density (PSD) and the fit of the aperiodic component of the regions with the maximum t-value in each of the comparisons performed. Cortical maps are thresholded at *p* < 0.05 after FDR correction.

**Figure 3.**
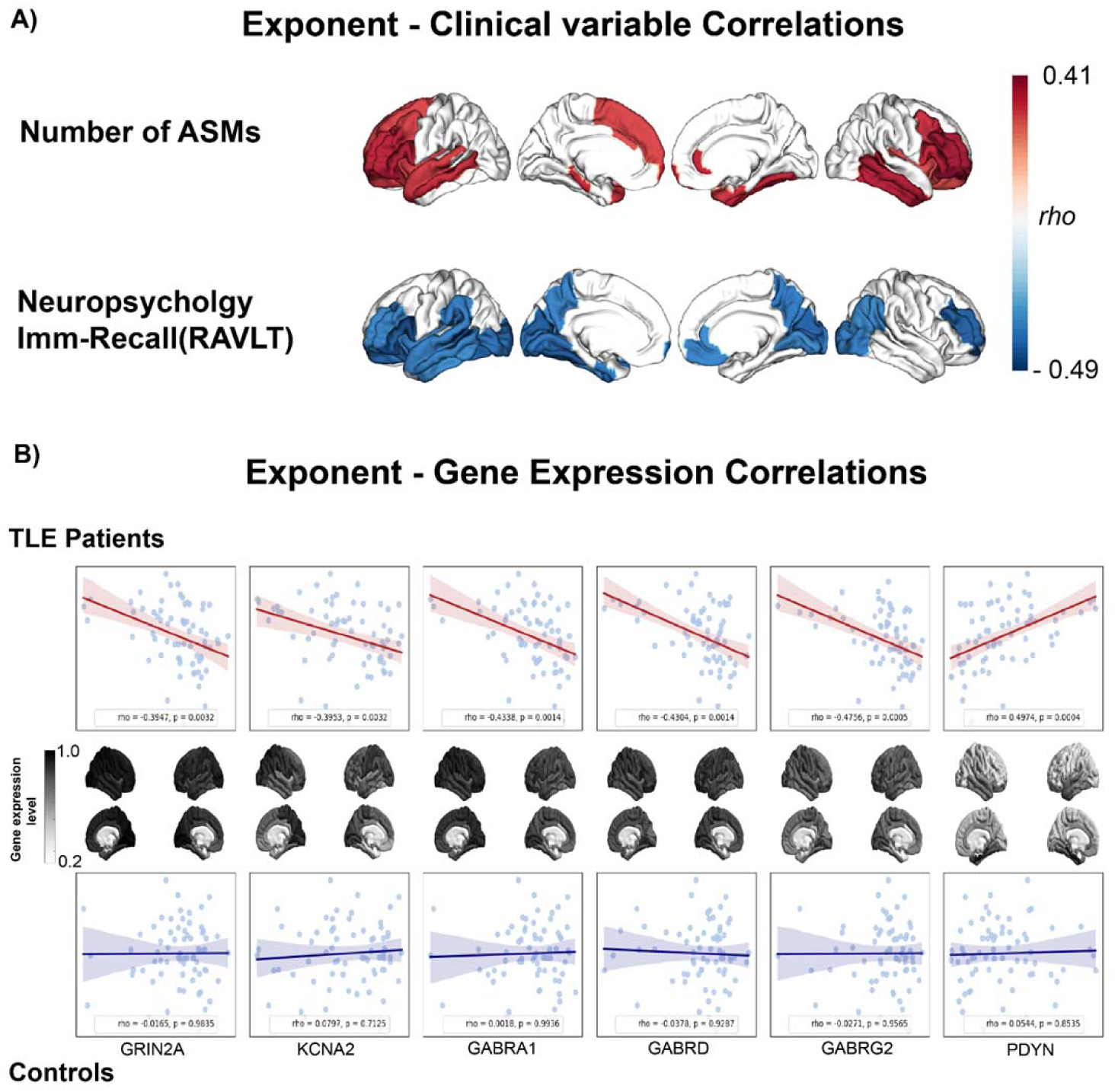
Exponent correlation with clinical and transcriptomics variables. Panel A shows the spatial distribution on the cortex of the node-wise correlation between region exponent and the number of antiseizure medications (ASMs) and the short term verbal memory functioning measured by the immediate recall (Imm-Recall) of the Rey Auditory Verbal Learning Test (RAVLT). Panel B displays the exponent-gene expression correlation both for patients with TLE (upper line) and the control group (lower line). The brain plots in black and white represent the node-wise cortical expression of the corresponding gene.

## 5 Limitations

We cannot rule out that the exponent of the PSD captures other signal properties beyond E/I balance (e.g., criticality; He et al., 2014). The largest part of the literature investigating the PSD features in epilepsy has seldom disentangled the periodic from the aperiodic component. Patients with TLE are known to have increased slow activity ^102,103^. The mechanisms sustaining these phenomena are manyfold and they involve structural damage ^104^, cortical plasticity ^105,106^ and intracranial epileptiform abnormalities which appear as slow activity on the surface due to the conductance properties of skin and skull ^103,107,108^. Simultaneous invasive and non-invasive recordings may be beneficial to parametrize the influence of these components on the exponent-derived estimation of E/I ^11^.

## 6 Conclusions

Our results reinforce the aperiodic exponent as a functional measure of E/I of the intrinsic functional organization on a large scale network in patients with TLE, displaying sensitivity to the lateralization of the epileptogenic hemisphere. The correlational findings support the functional relevance of the E/I assessed based on the aperiodic exponent for cognition, with a dependence on antiseizure medication. The syndrome-specific functional-transcriptomic signatures could provide information on the altered mechanisms underlying the E/I balance deregulation implied in the etiopathogenesis epilepsy. In this study we evidenced the potential usage of a cost-efficient and non-invasive measure of the E/I balance. Further studies may highlight the potential value of integrating the simple and informative measures presented here, in the assessment of TLE patients.

## Data availability statement

The data that support the findings of this study are available on request to the corresponding author. The raw data are not publicly available due to privacy or ethical restrictions. All the scripts are available at the following github page:

https://github.com/simone-cuozzo/FOOOF_epilepsy

