## Supplementary Materials for "Excitation/Inhibition balance relates to cognitive function and gene expression in Temporal Lobe Epilepsy: an hdEEG assessment with aperiodic exponent"

| Studied Genes | Area of Interest |
| --- | --- |
| LGI1 | Temporal Epilepsy |
| CNTNAP2 | Cortical dysplasia |
| GRIN2A | Focal epilepsy |
| SCN1A | Sodium channels |
| SCN1B | Sodium channels |
| SCN2A | Sodium channels |
| KCNA2 | Potassium channels |
| KCNB1 | Potassium channels |
| KCNC1 | Potassium channels |
| KCNMA1 | Potassium channels |
| KCNQ3 | Potassium channels |
| KCNT1 | Potassium channels |
| GABRA1 | Gaba receptor |
| GABRB1 | Gaba receptor |
| GABRB2 | Gaba receptor |
| GABRB3 | Gaba receptor |
| GABRD | Gaba receptor |
| GABRG2 | Gaba receptor |
| GRIN2B | NMDA receptor |
| GRIN2D | NMDA receptor |
| GRINA | NMDA receptor |

| | |
|---|---|
| PDYN | Seizure suppression |

**Supplementary Table 1.** The present table enlists the gene and the corresponding function, selected for the gene expression-exponent correlation.

| ALL PATIENTS vs. CONTROLS | | |
|---|---|---|
| **Regions** | **t-values** | **p-values** |
| bankssts R | 2,7287 | 0,0176 |
| caudalanteriorcingulate R | 3,1853 | 0,0064 |
| cuneus L | 2,9242 | 0,0119 |
| cuneus R | 3,6811 | 0,0021 |
| entorhinal L | 3,9091 | 0,0014 |
| entorhinal R | 4,2011 | 0,0007 |
| frontalpole L | 3,4686 | 0,0036 |
| frontalpole R | 3,3505 | 0,0043 |
| fusiform L | 2,9655 | 0,0119 |
| inferiortemporal L | 2,3066 | 0,0448 |
| inferiortemporal R | 3,2686 | 0,0056 |
| insula L | 3,0884 | 0,0083 |
| insula R | 2,491 | 0,0283 |
| isthmuscingulate R | 3,7996 | 0,0017 |
| lateraloccipital L | 4,3179 | 0,0007 |
| lateraloccipital R | 4,1457 | 0,0007 |
| lingual L | 2,8372 | 0,0138 |
| medialorbitofrontal L | 6,2482 | 5.5e-07 |
| medialorbitofrontal R | 4,3958 | 0,0005 |
| middletemporal L | 3,0668 | 0,0089 |
| middletemporal R | 3,7958 | 0,0017 |
| paracentral L | 2,8148 | 0,0138 |
| paracentral R | 2,85 | 0,0134 |
| parahippocampal L | 2,6845 | 0,0196 |

| | | |
|---|---|---|
| parsopercularis L | 3,3897 | 0,0042 |
| parsorbitalis L | 3,2909 | 0,005 |
| parsorbitalis R | 3,7517 | 0,0017 |
| parstriangularis L | 3,4789 | 0,0036 |
| parstriangularis R | 3,0788 | 0,0083 |
| precuneus L | 2,6441 | 0,0196 |
| precuneus R | 2,6955 | 0,0181 |
| rostralanteriorcingulate L | 4,1638 | 0,0007 |
| rostralanteriorcingulate R | 3,4396 | 0,0037 |
| superiortemporal L | 2,5153 | 0,028 |
| temporalpole L | 6,3984 | 5.5e-07 |
| temporalpole R | 5,0209 | 5.4e-05 |
| transversetemporal L | 2,5597 | 0,025 |

**Supplementary Table 2.** The present table enlists the *t*- and the FDR corrected *p-values* for each significant region in the comparison of the aperiodic exponent between the whole group of patients with epilepsy and healthy controls

| LEFT TLE vs. CONTROLS | | |
|---|---|---|
| **Regions** | **t-values** | **p-values** |
| bankssts L | 2,6589 | 0,0348 |
| caudalanteriorcingulate R | 2,649 | 0,0348 |
| cuneus L | 2,6527 | 0,0348 |
| cuneus R | 2,8388 | 0,0288 |
| entorhinal L | 3,9742 | 0,0033 |
| entorhinal R | 3,0777 | 0,0167 |
| fusiform L | 2,8529 | 0,0288 |
| insula L | 3,6415 | 0,0062 |
| isthmuscingulate R | 3,3049 | 0,0118 |
| lateraloccipital L | 4,0948 | 0,0027 |
| lateraloccipital R | 2,5161 | 0,0437 |

| | | |
|---|---|---|
| lingual L | 2,7197 | 0,0337 |
| medialorbitofrontal L | 5,1364 | 0,0001 |
| medialorbitofrontal R | 3,2228 | 0,0131 |
| middletemporal L | 3,5648 | 0,0062 |
| parahippocampal L | 2,5468 | 0,0411 |
| parsopercularis L | 3,2769 | 0,0118 |
| parsorbitalis L | 2,7445 | 0,0337 |
| parstriangularis L | 3,5696 | 0,0062 |
| rostralanteriorcingulate L | 3,0908 | 0,0167 |
| superiortemporal L | 2,6893 | 0,0345 |
| temporalpole L | 5,2419 | 0,0001 |
| temporalpole R | 3,5982 | 0,0062 |

**Supplementary Table 3.** The present table enlists the *t*- and the FDR corrected *p-values* for each significant region in the comparison of the aperiodic exponent between the group of patients with left temporal lobe epilepsy (Left-TLE) and healthy controls

| RIGHT TLE | | |
|---|---|---|
| **Regions** | **t-values** | **p-values** |
| temporalpole L | 3,9774 | 0,0224 |

**Supplementary Table 4.** The present table enlists the *t*- and the FDR corrected *p-values* for each significant region in the comparison of the aperiodic exponent between the group of patients with right temporal lobe epilepsy (Right-TLE) and healthy controls

| BILATERAL TLE | | |
|---|---|---|
| **Regions** | **t-values** | **p-values** |
| bankssts L | 2,5844 | 0,0236 |
| bankssts R | 4,1348 | 0,0013 |
| caudalanteriorcingulate L | 2,5594 | 0,0271 |

| | | |
|---|---|---|
| caudalanteriorcingulate R | 2,9396 | 0,0132 |
| caudalmiddlefrontal L | 2,4715 | 0,0303 |
| caudalmiddlefrontal R | 2,4733 | 0,0298 |
| cuneus L | 2,4796 | 0,0298 |
| cuneus R | 2,8236 | 0,0173 |
| entorhinal L | 4,1744 | 0,0013 |
| entorhinal R | 5,2479 | 0,0002 |
| frontalpole L | 4,0355 | 0,0017 |
| frontalpole R | 4,0217 | 0,0017 |
| fusiform L | 3,8975 | 0,0013 |
| inferiortemporal L | 2,4883 | 0,0277 |
| inferiortemporal R | 4,7822 | 0,0005 |
| insula L | 3,1311 | 0,0081 |
| insula R | 4,0093 | 0,0013 |
| isthmuscingulate R | 4,1153 | 0,0013 |
| lateraloccipital L | 3,1139 | 0,0089 |
| lateraloccipital R | 4,38 | 0,0012 |
| lateralorbitofrontal L | 2,6286 | 0,0235 |
| lateralorbitofrontal R | 3,0227 | 0,0111 |
| medialorbitofrontal L | 5,3315 | 0,0003 |
| medialorbitofrontal R | 4,565 | 0,001 |
| middletemporal L | 3,2124 | 0,0075 |
| middletemporal R | 3,904 | 0,0019 |
| paracentral L | 2,5725 | 0,0271 |
| paracentral R | 2,7004 | 0,0229 |
| parahippocampal L | 3,4528 | 0,0037 |
| parahippocampal R | 2,4161 | 0,033 |
| parsopercularis L | 3,3579 | 0,0068 |
| parsopercularis R | 3,0794 | 0,0082 |
| parsorbitalis L | 3,2172 | 0,0081 |
| parsorbitalis R | 5,4299 | 0,0002 |
| parstriangularis L | 3,8384 | 0,0019 |

| | | |
|---|---|---|
| parstriangularis R | 4,2053 | 0,0013 |
| precuneus R | 2,6622 | 0,0235 |
| rostralanteriorcingulate L | 4,0842 | 0,0014 |
| rostralanteriorcingulate R | 3,7876 | 0,0027 |
| rostralmiddlefrontal L | 2,8046 | 0,0173 |
| rostralmiddlefrontal R | 3,281 | 0,0076 |
| superiorfrontal L | 2,2703 | 0,0451 |
| superiortemporal L | 3,9273 | 0,0013 |
| superiortemporal R | 3,8424 | 0,0017 |
| temporalpole L | 4,809 | 0,0006 |
| temporalpole R | 4,4478 | 0,0013 |
| transversetemporal L | 3,1626 | 0,008 |
| transversetemporal R | 3,1949 | 0,008 |

**Supplementary Table 5.** The present table enlists the *t*- and the FDR corrected *p-values* for each significant region in the comparison of the aperiodic exponent between the group of patients with bilateral temporal lobe epilepsy (BTLE) and healthy controls

| **Exponent – number of ASMs Spearman Correlation** | | |
|---|---|---|
| **Regions** | **rho** | **p-values** |
| bankssts R | 0,353 | 0,0258 |
| caudalanteriorcingulate L | 0,2839 | 0,0467 |
| caudalmiddlefrontal L | 0,295 | 0,0418 |
| caudalmiddlefrontal R | 0,3146 | 0,0329 |
| entorhinal R | 0,3139 | 0,0329 |
| frontalpole L | 0,3529 | 0,0258 |
| frontalpole R | 0,3432 | 0,027 |
| fusiform R | 0,3359 | 0,027 |
| inferiortemporal R | 0,3444 | 0,027 |
| insula L | 0,2992 | 0,0394 |
| insula R | 0,286 | 0,0463 |
| lateralorbitofrontal L | 0,3061 | 0,0381 |

| | | |
|---|---|---|
| lateralorbitofrontal R | 0,2818 | 0,0473 |
| middletemporal L | 0,3189 | 0,0329 |
| middletemporal R | 0,3545 | 0,0258 |
| parahippocampal L | 0,3293 | 0,0277 |
| parsopercularis L | 0,356 | 0,0258 |
| parsopercularis R | 0,3326 | 0,027 |
| parsorbitalis L | 0,3831 | 0,0258 |
| parsorbitalis R | 0,4098 | 0,0258 |
| parstriangularis L | 0,3527 | 0,0258 |
| parstriangularis R | 0,3584 | 0,0258 |
| rostralanteriorcingulate R | 0,3335 | 0,027 |
| rostralmiddlefrontal L | 0,3392 | 0,027 |
| rostralmiddlefrontal R | 0,3664 | 0,0258 |
| superiorfrontal L | 0,2934 | 0,0418 |
| superiortemporal L | 0,3017 | 0,0387 |
| temporalpole L | 0,3164 | 0,0329 |
| temporalpole R | 0,302 | 0,0387 |
| transversetemporal R | 0,2858 | 0,0463 |

**Supplementary Table 6.** The present table enlists the *rho* and the FDR corrected *p-values* for each significant region in the node-wise correlation in the patients group between the exponent value and the number of antiseizure medications.

| Exponent – Imm-recall RAVLT Spearman correlation | | |
|---|---|---|
| **Regions** | **rho** | **p-values** |
| bankssts L | -0,4696 | 0,0207 |
| cuneus L | -0,4444 | 0,0207 |
| cuneus R | -0,4319 | 0,0207 |
| entorhinal L | -0,3938 | 0,0347 |
| frontalpole L | -0,4428 | 0,0207 |
| fusiform L | -0,4405 | 0,0207 |
| inferiorparietal R | -0,3589 | 0,0406 |

| | | |
|---|---|---|
| inferiortemporal L | -0,3549 | 0,0407 |
| insula L | -0,3747 | 0,0372 |
| lateraloccipital L | -0,4043 | 0,0347 |
| lateraloccipital R | -0,3851 | 0,036 |
| lateralorbitofrontal L | -0,3925 | 0,0347 |
| lingual L | -0,3934 | 0,0347 |
| lingual R | -0,3566 | 0,0407 |
| medialorbitofrontal R | -0,3725 | 0,0372 |
| middletemporal L | -0,3677 | 0,0372 |
| parsopercularis L | -0,4796 | 0,0207 |
| parsorbitalis L | -0,4945 | 0,0207 |
| parstriangularis L | -0,4419 | 0,0207 |
| pericalcarine L | -0,394 | 0,0347 |
| pericalcarine R | -0,4347 | 0,0207 |
| precuneus L | -0,3737 | 0,0372 |
| precuneus R | -0,3877 | 0,036 |
| rostralanteriorcingulate R | -0,3588 | 0,0406 |
| rostralmiddlefrontal L | -0,3699 | 0,0372 |
| rostralmiddlefrontal R | -0,4327 | 0,0207 |
| superiortemporal L | -0,367 | 0,0372 |
| supramarginal L | -0,3793 | 0,0372 |

**Supplementary Table 7.** The present table enlists the *rho* and the FDR corrected *p-values* for each significant region in the node-wise correlation in the patients group between the exponent value and the immediate recall value of the Rey Auditory Verbal Learning Test.
